# Evolution of conserved noncoding sequences in *Arabidopsis thaliana*

**DOI:** 10.1101/727669

**Authors:** Alan E. Yocca, Zefu Lu, Robert J. Schmitz, Michael Freeling, Patrick P. Edger

**Affiliations:** Department of Plant Biology, Michigan State University, 612 Wilson Rd. East Lansing MI 48823; Department of Horticulture, Michigan State University, 1066 Bogue St. East Lansing, MI 48824; Department of Genetics, University of Georgia, 120 Green Street, Athens, GA 30602-7223; Department of Plant and Microbial Biology, University of California, 111 Koshland Hall, Berkeley, CA 94720; Ecology, Evolutionary Biology and Behavior, Michigan State University, East Lansing, MI, USA 48824

## Abstract

Recent pangenome studies have revealed a large fraction of the gene content within a species exhibits presence-absence variation (PAV). However, coding regions alone provide an incomplete assessment of functional genomic sequence variation at the species level. Little to no attention has been paid to noncoding regulatory regions in pangenome studies, though these sequences directly modulate gene expression and phenotype. To uncover regulatory genetic variation, we generated chromosome-scale genome assemblies for thirty *Arabidopsis thaliana* accessions from multiple distinct habitats and characterized species level variation in Conserved Noncoding Sequences (CNS). Our analyses uncovered not only evidence for PAV and positional variation (PosV) but that diversity in CNS is non-random, with variants shared across different accessions. Using evolutionary analyses and chromatin accessibility data, we provide further evidence supporting roles for conserved and variable CNS in gene regulation. Characterizing species-level diversity in all functional genomic sequences may later uncover previously unknown mechanistic links between genotype and phenotype.

## Introduction

Conserved noncoding DNA remains a highly understudied class of functional genomic features compared to protein-coding genes. Previous comparative genomic analyses in plants have identified stretches, generally 15-150 base pairs (bp) long (Fig S1), of noncoding regions that are positionally-conserved with identical (or near identical) sequence across distantly related species (1–4). These sequences, commonly referred to as Conserved Noncoding Sequences (CNS), are regions in the genome displaying much higher similarity across different taxa than expected by chance. Background mutation and genetic drift purges non-functional sequences over long evolutionary distances. Therefore, sequence conservation above expectation implies purifying selection actively conserves these CNS. Indeed, Williamson et al. (5) discovered elevated signatures of purifying selection in CNS regions compared to other classes of noncoding DNA in *Capsella grandiflora*. Previous studies demonstrated CNS contain transcription factor binding sites (TFBS) (2, 6, 7).

TFBS are typically 6-12 base-pair (bp) long (8). CNS can exceed this length, as they are thought to consist of arrays of TFBS capable of recruiting independent or cooperative transcriptional protein complexes. The length of CNS enables high confidence identification of orthologous cis-regulatory elements in comparator genomes. Querying genomes for TFBS alone results in a high false positive rate, as there are >30,000 expected occurrences of a given six bp sequence expected by chance even in the relatively small (~135 Mb) *Arabidopsis thaliana* genome. In contrast, there is less than one expected random occurrence of the shortest CNS (15bp). TFBS colocalize with accessible chromatin in mammalian genomes, as do CNS as demonstrated previously in plants (4, 9–14).

Alexandre and coworkers previously investigated variation in signatures of accessible chromatin and sequence diversity of differentially accessible regions across five diverse *A. thaliana* accessions (15). They discovered ~15% of accessible chromatin regions differed across the five accessions, with a minority of those sites displaying sequence divergence. Mapping data from non-reference genotypes to a reference genome may result in incorrectly assigning reads to the reference (16). By assembling separate genomes for each accession, we mitigate this reference mapping bias. We add to the findings of Alexandre and coworkers by characterizing sequence diversity directly on a larger panel of thirty accessions, focused on a CNS set consisting of more than 3Mb of sequence, and discover significant relationships between variable sequence and regions of accessible chromatin. Genes experience a broad spectrum of selective forces potentially resulting in strong conservation (i.e. resisting deletion) (17) or active removal from certain genomes (18). Certain gene families are known to exhibit high birth-death dynamics because ancestral genes are lost from the lineage, whereas other gene families are relatively stable in size (19, 20). Thus, some genes are present in all eukaryotes, whereas others may be lineage specific (21). Equivalently, a subset of CNS identified across Brassicaceae (1) are identifiable across all surveyed angiosperms including *Amborella* (2) whereas others are uniquely shared by only a subset of Brassicaceae.

Previous pangenome studies aimed to capture presence-absence variation in transcribed regions to characterize the core and dispensable gene content, however, these studies often focus on *de novo* assembly of only the non-reference gene space (22–25). These studies consistently find core genes (those present across most individuals within a species) are enriched in essential cellular processes, whereas dispensable genes often display higher mutation rates and are biased towards adaptive processes (e.g. response to the environment). We hypothesize dispensable CNS follow patterns observed for dispensable coding regions such as representing a pool of sequences contributing to adaptive processes and potentially important agronomic traits.

Though tens of thousands of CNS have already been identified in plant genomes, these comparisons are often performed between single representatives of select distantly related species. To our knowledge, the variation in CNS content across the genome of multiple individuals within a single species has never been addressed in plants. Here, we assembled chromosome-scale genomes for thirty *A. thaliana* accessions and leveraged one of the largest annotated CNS datasets (1), to investigate the levels and patterns of intraspecific variation of CNS and the impact of this variation on gene expression in *A. thaliana*.

## Results

### What proportion of CNS vary within a species?

CNS are typically identified through whole genome comparisons of single representative genomes of different species spanning various phylogenetic distances. Therefore, the variation of these sequences at the species level remains poorly understood, especially in plants. We investigate two main types of variation in CNS structure across multiple *A. thaliana* accessions: presence-absence variation (PAV) and positional variation (PosV). We define PAV CNS as those present in the reference accession (Col-0), but absent in at least one other accession. PosV CNS are those which exist in a different locus in an accession relative to its position in Col-0. A model of intraspecific CNS variation is shown in Fig 1A.

**Figure 1:**
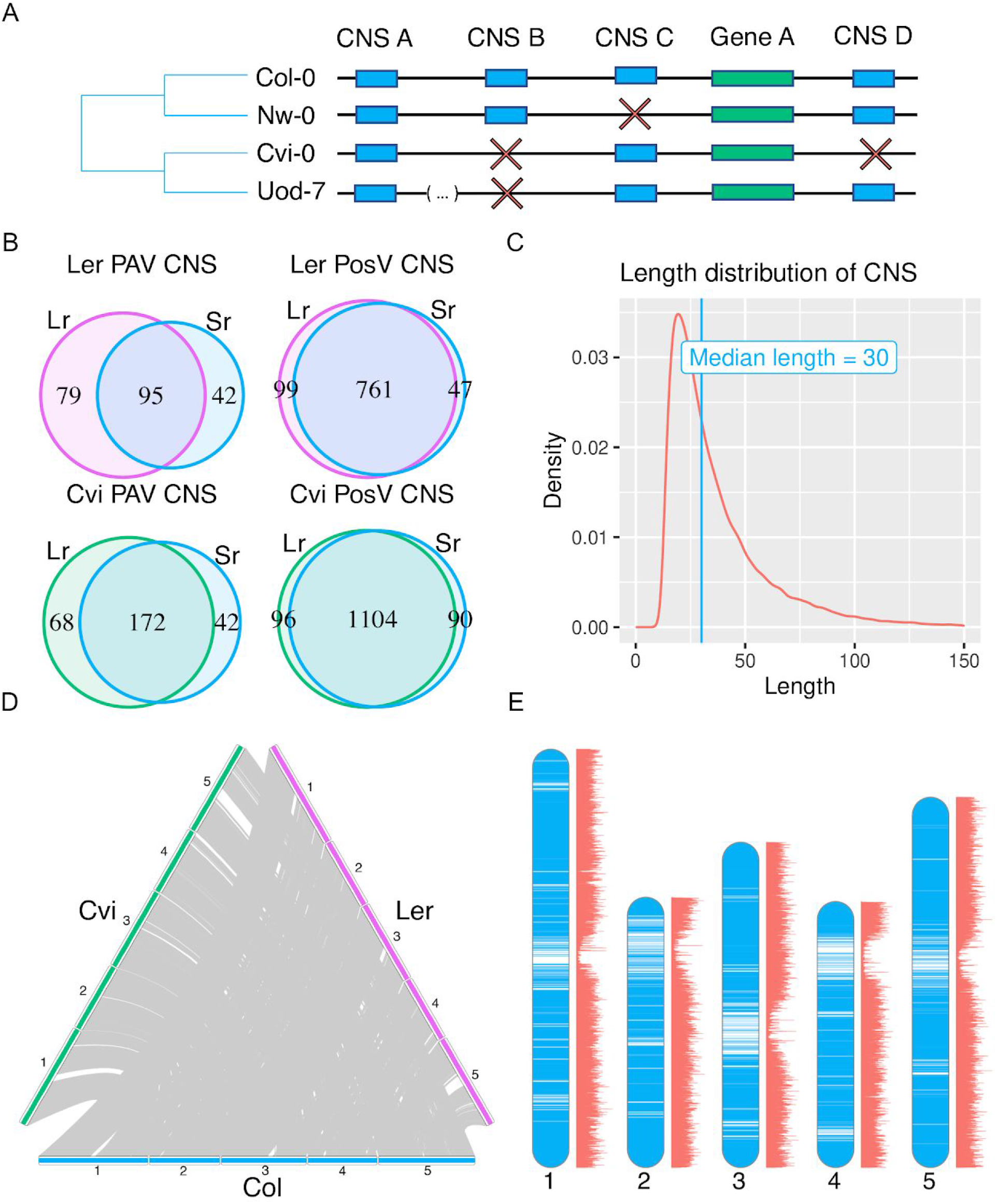
(A) A model depicting intraspecific variation in CNS content. Blue blocks represent conserved non-coding sequences (CNS) and green blocks represent genes. CNS may occur upstream or downstream genes. Red “X” characters depict the failure to identify a CNS in an *A. thaliana* accession in the position at which we find the CNS in the reference accession Col-0. CNS A in accession Uod-7 is found at a location other than where it is found in Col-0 as demarcated by the genomic position break. Therefore, in accession Uod-7, CNS A displays PosV. (B) The length distribution in base-pairs of the 62,916 CNS studied here showing a minimum sequence length of fifteen base-pairs, and a median of thirty base-pairs. (C) Venn diagrams comparing the CNS variation observed in genetically identical accessions between publically available long-read (Lr) genome assemblies and those generated with short-reads (Sr) in this study. (D) A three-way synteny diagram showing collinear CNS structure (i.e. not displaying PAV nor PosV) between two accessions, Ler-0 (purple) and Cvi-0 (green), and the reference accession Col-0 (blue). (E) Location of collinear CNS in the Ler-0 long-read genome assembly relative the the reference accession Col-0 are shown in blue overlaying the karyotype. ATAC-seq read depth is shown adjacent to each chromosome in red with depth increasing along the horizontal axis.

Importantly, we only investigate CNS present in the *A. thaliana* reference accession Col-0 identified by Haudry, *et al* (1). These elements were identified through whole genome alignments of nine phylogenetically informative taxa within Brassicaceae resulting in a set of >60,000 CNS used in this study whose length distribution is shown in Fig 1B. Regions were classified as CNS if they exhibited strong conservation across most investigated taxa. Importantly, these elements also displayed variation across Brassicaceae, but were broadly present across taxa. As the Col-0 accession was the reference genotype for CNS identification, we do not consider CNS present in non-reference accessions which are absent in Col-0.

To investigate species level CNS variation, we queried seven recently available long-read sequencing *de novo* genome assemblies for various *A. thaliana* accessions (Table S5; Table S6; 26). As the set of query CNS were characterized for their presence in multiple different species across Brassicaceae spanning ~32 million years of evolution (27), low proportions of variation were expected. Indeed the vast majority of CNS are conserved as demonstrated in Fig 1C. Of the 62,916 CNS investigated, we find an average of 209 (0.33%) and 951 (1.5%) CNS exhibit PAV and PosV respectively per accession. Querying seven accessions may be insufficient to capture a large extent of the natural genetic variation occurring in these sequences as *A. thaliana* has a global distribution and at least nine definable genetic admixture groups (28).

Therefore, we assembled the genomes of thirty *A. thaliana* accessions using a hybrid reference and *de novo* method (Supplementary Methods). This included assemblies for two accessions for which a long-read assembly was available for direct comparison (Ler-0 & Cvi-0). We find appreciable overlap in the PAV and PosV CNS identified between our assemblies and the long-read assemblies (Table S5; Table S6; Fig 1D). Of the 62,916 CNS analyzed, we find an average of 163 (0.26%) and 910 (1.4%) CNS exhibit PAV and PosV respectively per accession. These estimates are in line with those obtained using the long-read genome assemblies. Given the large number of CNS in the query set (62,916), this represents a definable class of sequence (>1,000 sequences per accession) with observable variation patterns, and these we examine further. The subsequent analyses were performed on the larger set of thirty assemblies we generated.

Throughout the manuscript, CNS exhibiting PAV in at least one accession will be referred to as PAV CNS. A similar syntax will follow for CNS showing PosV in at least one accession. CNS in either of the aforementioned classes will be referred to as variable CNS, whilst those showing no variation are referred to as collinear CNS.

### Is CNS variation shared among accessions?

CNS variation is highly shared among accessions. If PAV and PosV CNS occurred independently in each accession, we expect 4,567 and 21,699 different CNS to be lost and positionally variable respectively in at least a single accession (Supplementary Methods). In contrast to random expectation, we only observe 1,524 and 4,801 distinct CNS lost and positionally variable respectively (Fig S2; Fig S3). Therefore, PAV and PosV CNS events are likely non independent among individuals. This indicates variable CNS are often variable in more than a single accession. There is little overlap of PAV and PosV CNS. There are 118 CNS absent in at least one accession and positionally variable in at least one other accession (<10% of either set). This is not significantly different than expected by chance (hypergeometric test p-value = 0.4227782).

Random subsampling of CNS analyzed in the thirty accessions assembled indicate the majority of the natural common CNS variation is likely captured and is sufficient to investigate the functional consequences of this variation (Fig S2; Fig S3). Furthermore, there is strong observed overlap in variable CNS across accessions (Fig S4; Fig S5), indicating there may be subclasses of CNS which are more likely to exhibit variation than others. This phenomenon is similar to certain classes of gene families that often display copy-number variation (19, 29, 30).

### How does CNS variation compare to gene content variation?

Previous studies devoted major efforts to pan-genome analysis while assessing species level diversity in gene content and structural variants (22–25). Most of these studies often do not fully assemble genomes for each individual of the species. Rather, they only assemble the sequence not present in the reference. It is challenging to identify positional conservation and rearrangements in non-reference individuals using these approaches. However, some previous pangenome studies (e.g. *Brachypodium*; 22) have assembled full genomes but focused on only coding gene content variation. Our approach uses a hybrid reference guided and *de novo* assembly approach to obtain chromosome-scale sequences for each individual accession. This permits the analysis of PAV and positional variation (PosV) of both CNS and transcriptional unit content.

Our analyses revealed that CNS variation occurs at a much lower rate than genic presence-absence variation (Fig S6). However, the calculated rate of gene PAV relative to the reference accession Col-0 may be overestimated due to greater annotation quality for the reference. This might imply purifying selection acts more strongly on noncoding regulatory regions than protein-coding genes. However, the set of noncoding regions investigated in this study are also present throughout Brassicaceae, biasing our annotations to CNS likely experiencing greater levels of purifying selection. The true rate of variation in functional noncoding regions may only be identified through complete annotation of functional *cis*-regulatory regions, a difficult feat relative to coding region annotation. Thus, it is imperative that future efforts identify species-specific CNS to assess the full scope of regulatory variation that exists at the species level.

### What is the length distribution of variable CNS?

The distribution of the lengths of CNS was investigated (Fig S7). CNS retaining their syntenic position in every accession (collinear CNS) have a length distribution similar to that of all CNS in the reference accession. PAV CNS on average have a longer length (in base-pairs) than collinear CNS (collinear average = 39.84, PAV average = 44.41, *KS* test p-value < 2.2e^−16^). The PAV CNS length distribution appears slightly bimodal (Fig S7). PosV CNS are much shorter on average than either collinear or PAV CNS (PosV average = 18.97).

### What is the distribution of CNS movement events?

PosV CNS are those found outside of their respective syntenic block in the reference genotype. In other words, their order among CNS is different in the accession compared to the Col-0 reference genome. We observe an apparent bias in PosV location towards gene-rich regions (KS test p-value < 2.2 × 10^−16^; Fig S8).

In addition to genomic positional changes, CNS distance to their proximate gene was investigated. Fig 2 compares the distance of CNS to their proximate genes across a few different classes of CNS. The largest concentration of CNS is intergenic and close to genes in the genome (37.17% of CNS +/− 500 bp of and between transcriptional start or termination sites). There is a reduction in the concentration of CNS around the nearest gene for PosV CNS, relative to CNS in accessions that retain their syntenic position relative to Col-0. Position relative to the proximate gene does not seem to predispose a CNS from exhibiting PAV in the global population.

**Figure 2:**
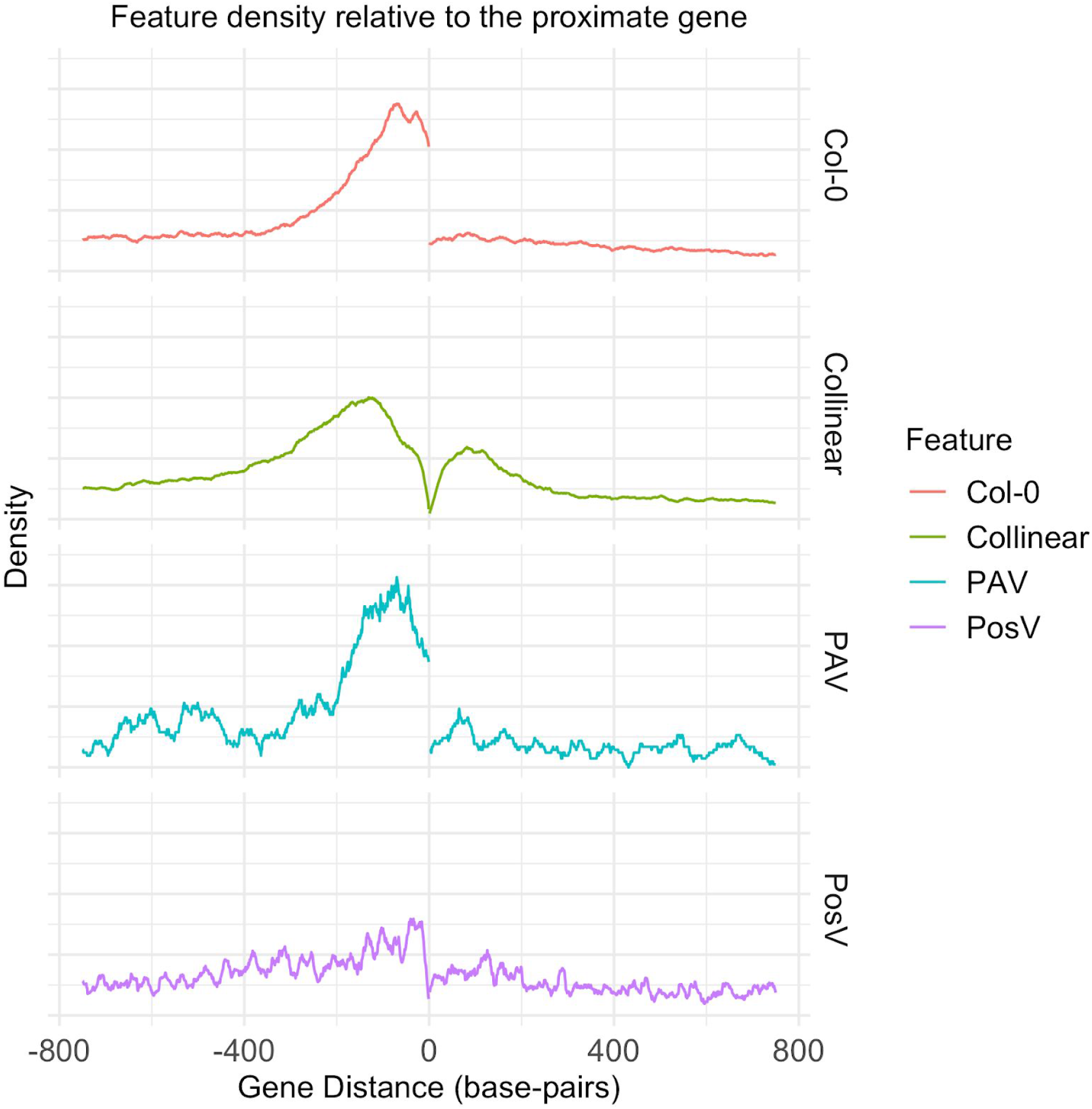
Distributions of different features relative to their proximate gene are shown, where the x-coordinate zero represents the location of the proximate gene. The top panel (gold) shows all CNS in the reference accession Col-0. The second panel (green) shows across all accessions the distribution of CNS that remain in the same syntenic position as the reference accession (Collinear CNS). The middle panel (blue) shows the position in the reference accession where PAV CNS are located, i.e. the position in Col-0 where CNS display PAV in accessions. The fourth row (purple) shows the position of PosV CNS across all accessions, i.e. the location to which these CNS “moved”. The final panel (red) shows the distribution of ATAC sequencing peaks across all accessions sampled.

### Does CNS structure align with stimuli experienced by different Arabidopsis accessions?

Principal Component Analysis (PCA) was performed to examine similarities in CNS variation across accessions. PCA was performed separately using PAV CNS and PosV CNS as input. The first two principal components (PCs) for PAV CNS explained 10.8% and 8.29% of the total variance. There are two distinct clusters of accessions defined by the first two PCs of this analysis for PAV CNS (Fig 3A). The first two PCs for PosV CNS explained 6.83% and 5.57% of the total variance. Two distinct clusters are apparent for PosV CNS, similar to PAV CNS (Fig 3C). Clustering accessions according to SNPs produced a topology similar to that constructed from PosV CNS information (Fig 3E). Jointly using PAV and PosV CNS as clustering information produced topology similar to using PosV CNS alone (Fig S9). We investigated whether clustering by CNS annotation aligns with bioclimatic variables obtained from WorldClim2 data (Fig 3; 33). With regard to ‘annual average temperatures’ (BIO1) at the location from where all accessions were collected, clustering according to CNS structure appears to separate accessions according to different distributions of environmental stimuli experienced by these accessions. The separation is much stronger for PCA clustering according to PosV CNS (Fig 3D) and SNPs (Fig 3F) than it is for PCA clustering according to PAV CNS (Fig 3B).

**Figure 3:**
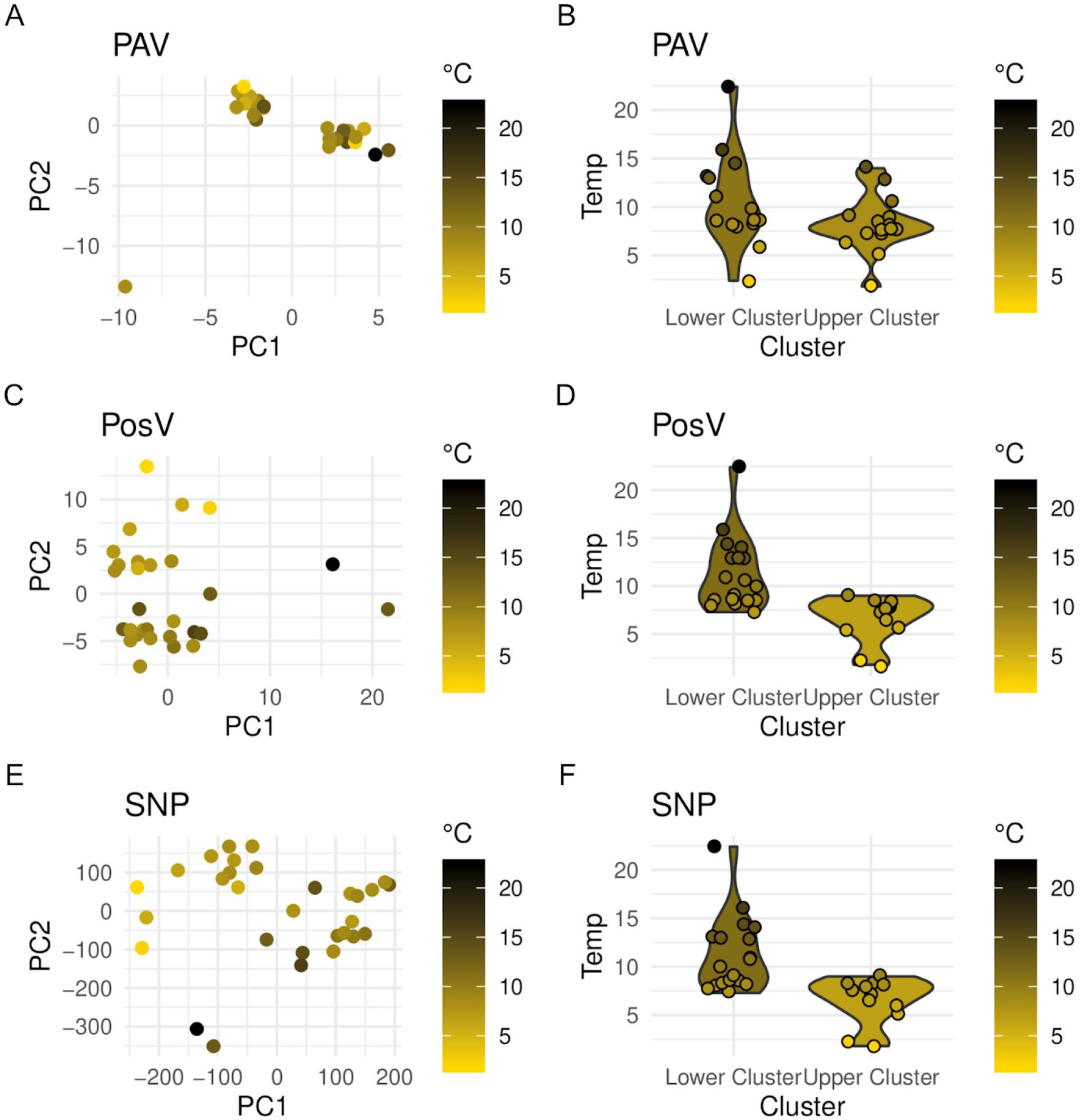
Principal Components Analysis (PCA) for each of the thirty accessions using either PAV CNS (A), PosV CNS (C), or single nucleotide polymorphism (SNPs; E) as information. Accessions are colored along a gradient according to the annual average temperature (BIO1) in degrees Celsius at the location from which the accession was collected. Violin plots (B, D and F) compare the distribution of the environmental variables between the different PCA clusters.

### Are variable CNS associated with accessible chromatin?

We performed Assay for Transposase-Accessible Chromatin sequencing (ATAC-seq) in leaf tissue for eighteen accessions, including the reference accession Col-0. This method identifies genomic regions accessible by a Tn5 transposase, and such regions of accessible chromatin are often associated with *cis*-regulatory DNA elements and transcription factor binding (13, 14, 34). We utilized a protocol which combines fluorescence-activated nuclei sorting and ATAC-seq (FANS-ATAC-seq; 12). As we hypothesize CNS are regulatory sequences, we expect that transcription factor binding will be enriched near regions of accessible chromatin. ATAC-seq reads were aligned to their respective genome, and peaks, regions of statistically enriched clusters of sequencing reads that are indicative of accessible chromatin, were identified. Collinear CNS demonstrated much stronger overlap with ATAC peaks than expected by chance (average fold-enrichment = 3.086; Table S1). Across all accessions, an average of 13.33% of CNS annotations overlapped chromatin accessible regions. Furthermore, we tested whether CNS which deviate from their position in the Col-0 accession (PosV CNS) still overlapped ATAC peaks. Nearly every accession displayed a significant enrichment of overlap between PosV CNS and ATAC peaks than expected by chance (Table S1). The average fold enrichment for PosV CNS across all accessions was 1.28. The average percent overlap was 5.59%, lower than observed for collinear CNS.

Strong evidence of CNS overlapping signatures of accessible chromatin has been reported previously (6, 9–11, 13). In each case, the set of CNS queried was different, with estimates ranging from 14% to 48% of CNS overlapping signatures of accessible chromatin. The percentage reported here, 16%, is in line with previous estimates. In Fig 4, we demonstrate an instance where CNS loss is associated with loss of accessible chromatin in a given accession. Additionally, we show a novel CNS insertion in an accession associated with an accession-specific chromatin accessible region. Fig 4C provides limited quantitative support of an association between accessibility and CNS presence or absence, with an increase in accessibility upstream genes associated with a greater number of CNS compared to the reference accession. Identification of lineage specific regulatory regions in future studies may further illuminate the quantitative relationship between CNS and accessible chromatin.

**Figure 4:**
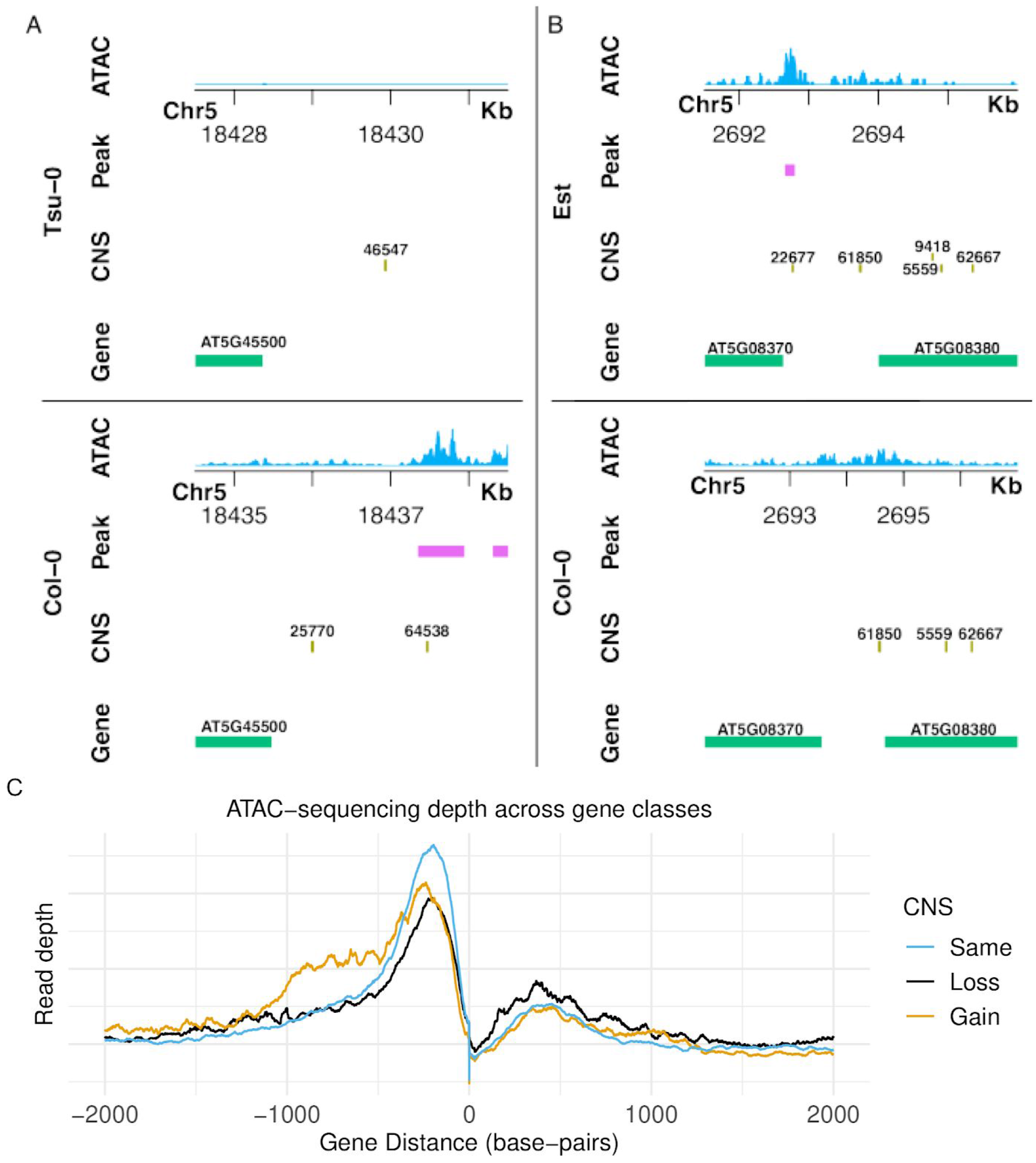
Genome browser tracks are shown for two different syntenic regions. (A) Shows loss of CNS 64538 and 25770 associated with loss of ATAC-seq peaks downstream the locus AT5G45500. Part (B) demonstrates a gain of CNS 22677 associated with the gain of a CNS peak upstream the locus AT5G08370. Coordinates are relative to each genome assembly. Thus orthologous sequences may have different coordinates due to insertions or deletions occurring in “earlier” coordinates. (C) ATAC-sequencing depth is shown surrounding orthologous genes with either the same (blue), less (black), or more (orange) CNS associated with them relative to the reference accession Col-0.

Though there is significant overlap between PosV CNS and accessible chromatin, the majority of PosV CNS do not overlap ATAC peaks. The true proportion of putatively functional PosV CNS is likely greater than observed here, potentially because the ATAC sequencing was performed at a single time point in a single organ under normal growing conditions. As CNS are hypothesized to perform regulatory functions, their binding partner may require distinct spatial-temporal context and/or environmental stimuli to activate. Given bulked tissue samples and homogenous environmental stimuli were sampled for the ATAC sequencing data, it is unlikely every PosV CNS which may exist in regions of accessible chromatin under different conditions or cell-types will be identified. In addition to responding to distinct environmental stimuli, regulatory functions of some CNS may also be cell/tissue/organ or developmental stage specific, further lending to their absence in regions of accessible chromatin observed here. Lastly, PosV CNS and ATAC peak overlap was lower than that of collinear CNS. PosV CNS may act as adaptive sequences, changing over shorter evolutionary distances, similar to certain classes of genes that exhibit higher transposition and duplication rates (20, 29, 35). They may be involved in specific stress responses and therefore may not demonstrate overlap with accessible chromatin in healthy leaves. However, future work is required to assess specifically whether PosV CNS exhibit these behaviors.

It should be noted that previous studies posited accessible region differences between *A. thaliana* cell types were primarily quantitative rather than qualitative (36). Perhaps PosV CNS not overlapping with accessible chromatin align with this trend and exhibit low signatures, rather than the absence, of accessible chromatin below our detection threshold. This is worth investigating in the future.

### Is CNS loss-and-gain associated with gene expression differences?

RNA-sequencing (RNA-seq) data were analyzed for four of the accessions investigated in this study to identify differentially expressed genes in leaf tissue. Each comparison was between an accession and Col-0 (37). Genes with a greater number of CNS associated with them in a given accession were more likely to be upregulated in that accession (Table 1). Genes with a lower number of CNS associated with them in a given accession were more likely to be downregulated in that accession. Genes without CNS changes relative to Col-0 were significantly underrepresented for differentially expressed genes. This demonstrates a significant association between changes in *cis*-regulatory sequence and divergent expression, a phenomenon also demonstrated across populations of stickleback fish (38). If the true ratio of activator binding sites to repressor binding sites were equal, we would expect no enrichment for differentially expressed genes for those gaining or losing CNS. Our results suggest CNS variation tilts towards a greater number or activity of *cis*-acting activator (enhancing expression) binding sites.

**Table 1:** Tests for over and under representation of differentially expressed genes in CNS with greater than (CNS Gain), less than (CNS Loss), or no change (No CNS Change) relative to Col-0. Numbers in parenthesis refer to the percent of the gene types which are up or down regulated.

| Gene Type | Average number of upregulated genes per accession (%) | P-value (Hypergeometric test) | Average number of downregulated genes per accession(%) | P-value (Hypergeometric test) |
| --- | --- | --- | --- | --- |
| CNS Gain | 208.5 (7.927) | $< 1 \times 10^{-10}$ | 76.5 (2.908) | $8.395 \times 10^{-06}$ |
| CNS Loss | 26.25 (2.449) | 0.093* | 121.75 (11.357) | $< 1 \times 10^{-10}$ |
| No CNS Change | 489.75 (2.212) | $< 1 \times 10^{-10}$ | 748.5 (3.380) | $< 1 \times 10^{-10}$ |

\*Did not pass Bonferroni correction p-value $0.01 / 6 = 0.00167$
Underrepresented
Overrepresented

### Are variable CNS enriched with certain binding motifs?

PAV and PosV CNS were searched for enriched motifs with the program HOMER (39). The set of all PAV and PosV CNS were tested separately. Motifs for the binding targets of several stress responsive transcription factor families were enriched. Specifically, in the set of PosV CNS sequences, the binding motifs of MYB113, C2H2, ABF3, HSF21, WRKY8, and CBF4 were enriched. For PAV CNS, WRKY50, RAV1, and Dof2 motifs were enriched. The global pattern for enriched motifs were for stress responsive elements. Given there are environmental differences experienced by different accessions (Fig 3), we hypothesize differences in regulatory patterns may govern an accession’s stress response.

One enriched motif in the set of PosV CNS is the target of HSFA4A. HSFA4A has been shown to regulate abiotic stress response pathways (40, 41) and has also shown accession-specific (Cvi-0) induction in response to biotic stresses (42). In the accession Cvi-0, the gene (AT4G18890) has lost 3 of 7 CNS that had been associated with it in the reference accession Col-0. A change in CNS structure may have resulted in rewiring of this transcription factor resulting in gene expression changes in certain environmental contexts.

### Are CNS changes associated with altered selective constraints?

As mentioned earlier, each CNS is associated with a gene in the Col-0 reference genome. Therefore, we can track the orthologous genes in each accession and determine if the genes in accessions which lose CNS exhibit signatures of positive or negative selection compared to those which have retained the CNS, including the Col-0 reference.

We assigned PosV CNS to their proximate gene to investigate changes in the CNS structure of orthologous genes across accessions. For example, we investigated any correlation between CNS gain or loss relative to Col-0 and Ka/Ks ratio (ratio of non-synonymous to synonymous substitution rate) for each gene in every accession. This revealed no clear correlation between Ka/Ks ratio and CNS count (Fig S10).

We were interested if a coding sequences’ association with a CNS in Col-0 altered the Ka/Ks pattern compared to those without an associated CNS. There was a statistically significant (KS test p-value < 1 × 10^−10^) difference between these two distributions indicating genes associated with a CNS in Col-0 generally display lower Ka/Ks values compared to genes with no associated CNS (all genes median Ka/Ks = 0.3853; CNS associated genes median Ka/Ks = 0.3549).

We were also interested if changes in CNS structure affect the Ka/Ks distribution, as calculated by comparing genes from each accession to its orthologous gene in the reference Col-0 accession. The Ka/Ks distribution for all gene pairs across all accessions is bimodal (Fig S11). The first peak is centered on 0, representing groups of identical shared alleles. The second peak occurs at a Ka/Ks ~ 0.45. The density of identical gene pairs is lower in orthologous pairs experiencing CNS change. There is, however, an increase in the density of orthologous gene pairs with a low Ka/Ks ratio (0.1 - 0.3). Given this observed CNS variation occurs at the species level, therefore a short evolutionary time frame, perhaps the change in CNS structure has already begun to exert positive selective pressure (evidenced by a higher Ka/Ks ratio).

The Ks distribution of different gene classes was investigated (Fig S12). The median Ks divergence of all orthologous gene pairs was 0.0040. The Ks distribution for all these gene pair groups showed a significant peak at 0, with a secondary peak at Ks ~ 0.003 followed by an exponential decay. The Ks distribution of orthologous gene pairs with less CNS in the accession relative to Col-0 (median Ks = 0.0169) and more CNS (median Ks = 0.0055) were observably different. Both distributions (CNS gain and loss) were shifted right. The CNS loss associated gene pair Ks distribution was the most shifted right.

### Are there relationships between CNS class and transposable elements or gene duplicates?

We investigated the proximity and density of transposable elements (TEs) for different classes of CNS. We identified putative TEs in each accession by searching for a set of previously annotated *A. thaliana* TEs within each genome using BLAST. Only matches at an e-value of lower than 1×10^−10^ were retained. Fig S13 demonstrates a clear bias in colocalization between PosV CNS and TEs relative to collinear CNS. Not only were PosV CNS on average closer to the nearest TE, but the density of TE matches both overlapping and within a 1,000 bp window was observably elevated in PosV CNS relative to collinear CNS. However, we observe elevated TE density for <25% of PosV CNS, indicating the majority of PosV CNS do not demonstrate elevated TE density relative to collinear CNS.

Lastly, we compared CNS content for different classes of duplicates (Fig S14). Considering only genes with a CNS associated with them, tandem duplicates had the fewest CNS associated with them (mean CNS count = 0.2442). This may be an artifact of CNS identification algorithms which struggle with tandem repeats. Genes without any duplicate in the genome (mean CNS count = 0.7064) had less CNS associated with them than genes with a duplicate pair dating back to the most recent whole genome duplication (At-alpha) shared by *A. thaliana* (mean CNS count = 0.9824) (27). This observation is consistent with previous studies; genes associated with CNS were more likely to be retained as duplicate pairs through diploidization potentially due to gene dosage constraints (43) or simply that these genes have long subfunctionalizable regulatory regions, or both explanations might be correct.

## Discussion

This study is, to our knowledge, the first genome-wide survey of CNS PAV and PosV at the species level in plants. The rate of variable CNS, while small compared to variable genes, is quite high in *A. thaliana*. However, the numbers reported here are likely underestimates of variable functional noncoding sequences given that our CNS set are heavily skewed towards those likely under stronger purifying selection. These positionally-conserved CNS were identified by aligning multiple Brassicaceae genomes spanning millions of years of evolution (1). Thus, new methods are needed to identify the full complement of functional regulatory sequences that are lineage and even species specific.

How is it that nearly 1,000 PosV CNS are at different loci in distinct accessions? We present two non-mutually exclusive hypotheses. First, we propose a *de novo* origin hypothesis. We find the distribution of PosV CNS to be noticeably shorter than the length distribution of all CNS (Fig S7). PosV CNS are often less than 20 bp in length. Therefore, perhaps the majority of the CNS sequence already exists in alternate loci, and only a few base-pair changes are needed to convert an existing background sequence to a CNS (Fig 5). A DNA sequence which is very similar to a binding motif may experience partial binding of a given transcription factor. This may be the selective pressure required to convert, or rather select for, beneficial mutations on the existing sequence to further strengthen that TF’s binding. Another possible explanation is that elevated mutation rates in recombination hotspots may contribute to the origin of PosV CNS. Future studies are needed to test whether PosV CNS sites are associated with regions of higher recombination. Second, the movement of regulatory elements may involve transposable elements (TEs) as shown previously (13, 44–46). Indeed, we observe strong bias with respect to the colocalization of PosV CNS and TEs relative to collinear CNS (Fig S13; Fig S15). These hypotheses are not mutually exclusive, and both may explain how PosV CNS arise at non reference locations.

**Figure 5:**
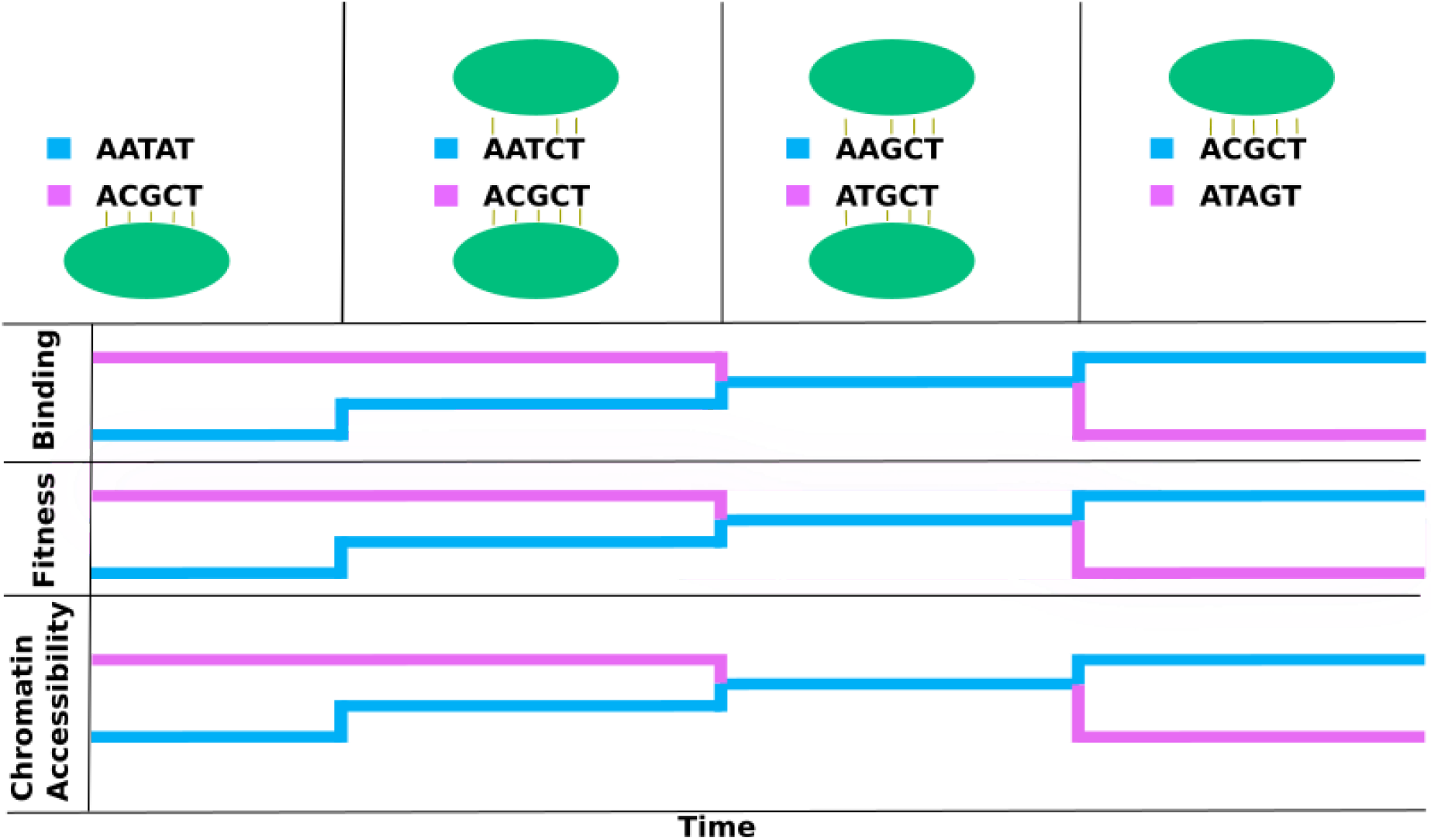
Hypothesized changes to transcription factor binding, fitness, and chromatin accessibility around two regions of DNA: the new PosV CNS location (top, blue), and the ancestral position as observed in Col-0 (bottom, pink). Each column in the first row depicts a snapshot of the state of the two CNS locations as time progresses along the x-axis. The hypothesized changes to TF binding, organism fitness, and chromatin accessibility align with the different time points.

Evolution of enhancer elements has been well studied in mammals (47, 48). These studies revealed thousands of lineage-specific enhancer elements have evolved across mammals and often occur in “ancient” DNA that is significantly under enriched for flanking repetitive elements. This suggests lineage-specific enhancer elements may arise through *de novo* origins via random mutations, in line with one of our hypotheses. Additionally, a few studies in *Drosophila* demonstrated *de novo* origins of TFBS (6bp-8bp) can occur on the order of 10^3^-10^6^ years under a model of neutral evolution (49, 50) which is within the divergence time (10^4^-10^5^ years) among *A. thaliana* accessions (28). Mustonen and Lassig (51) model binding site evolution in a manner we predict: CNS could occur *de novo*. According to their model, selective strength on random mutations depends upon the mutation’s effect on the binding strength of its associated transcription factor. Therefore, selection for partial transcription factor binding may drive sequence conversion from partial to full CNS sequence as shown in Fig 5.

Lastly, we provide evidence that positionally variable CNS retain significant associations with regions of accessible chromatin. Being open for business does not prove that business is actually being conducted, but functional genes must be open. Additional evidence for the function of PosV CNS, such as the effect of CNS change on gene expression, should be the focus of future studies in *A. thaliana*. This may need to involve genome editing of target CNS to assess its direct impact on gene expression and phenotype. We hypothesize PAV and shuffling of existing CNS at the population level serves as a mechanism to navigate the evolutionary landscape, providing more rapid alterations to the expression of genes; the consequent regulatory mutants would fuel fitness improvement.

## Data Availability

ATAC sequencing data are available under accession code PRJXXXXXX. Genome assemblies, gene annotations, and CNS annotations for each accession are deposited on Dryad (https://doi.org/10.5061/dryad.pzgmsbcfv).

## Author Contributions

All authors performed the research and/or analyzed data; A.E.Y and P.P.E. drafted the manuscript. All authors suggested experiments, reviewed and edited the manuscript.

## Acknowledgements

This work was supported by Michigan State University AgBioResearch and NSF-DEB 1737898 to P.P.E. A.E.Y. was supported by the College of Natural Science Recruiting Fellowship. M.F. supported by NSF Plant Genome Projects, R. Mosher, U. Arizona, PI, subaward UCB#041615-001. This work was supported in part by Michigan State University through computational resources provided by the Institute for Cyber-Enabled Research. R.J.S was supported by the NSF NSF IOS-1856627. R.J.S. is a Pew Scholar in the Biomedical Sciences, supported by The Pew Charitable Trusts.

## Methods

More details on all methods can be found in the Supplementary Methods. Genome assemblies, gene annotations, and CNS annotations will be made available on Dryad (https://doi.org/10.5061/dryad.pzgmsbcfv).

### Genome assembly

A separate genome assembly for each accession was generated using a hybrid reference-guided and *de novo* assembly method described in Supplementary Methods.

### Gene prediction

Genes were predicted for each accession independently using the MAKER pipeline.

### CNS annotation

CNS were annotated in each genome with the BLAST (52) program combined with stringent filtering. To increase the confidence in CNS annotations, all CNS below 15 base-pairs in length were dropped. After querying each ecotype’s genome with the final set of 62,916 CNS, the resulting hits were filtered to remove any match with a bit score lower than 28.2. Specifically, this corresponds to an exact 15 base-pair match. Hits were also dropped if they covered < 60% of the length of the CNS considered in that hit. Hits on separate chromosomes were also removed.

These steps were also followed for the reference genome to allow for accurate annotation of the CNS position in the reference. This annotation was used in comparison with that of each accession to determine CNS which exist outside of syntenic CNS blocks with a block size of 5. Synteny was determined using the MCScanX program (53).

### Identification of regions of accessible chromatin

Regions of accessible chromatin were identified using MACS2 (54).

### Ka/Ks calculation

Ka/Ks ratios were calculated using a series of custom script found on Github: https://github.com/Aeyocca/ka_ks_pipe. Briefly, syntenic orthologs were identified using JCVI Utilites Library (55) between each ecotype and the reference ecotype Col-0. The protein sequence of each pair was aligned using MUSCLE v3.8.31 (56). The protein alignment was converted to a coding sequence alignment using a modified version of PAL2NAL v14 (57) available on Github. Alignments were fed into PAML Version 4.9h (58) codeml function to calculate Ka/Ks values.

### PCA

Principal Components Analysis was performed with the prcomp() function in R version 3.5.0 (59). We decided not to scale variance, as CNS variability is not normally distributed.

### RNA-Sequencing

RNA sequencing data was taken for five accessions (Kn-0, Tsu-0, Ler-0, No-0, Col-0), from GSE30814 (37). Reads were mapped to their respective assembly using HISAT2/2.1.0 on default parameters (60). The resulting SAM file was converted to a BAM file using Picard Tools v2.18.1 (61). The BAM file was converted to a count matrix using StringTie/1.3.5 along with the accompanying script prepDE.py with minor modifications to handle variable gene names (62). The expression matrices were used to identify differentially expressed genes for each ecotype to the reference Col-0 using the R package DESeq2 (63).

### Motif Enrichment

Enriched motifs were identified using HOMER (39). Enrichment was performed for each ecotype separately as well as all ecotypes combined. For each set of ecotypes, enrichment was performed on PAV CNS and PosV CNS separately. The background set of sequences was set to a set of random sequences given the same composition of the query sequence using the scrambleFasta.pl script provided with HOMER.

### Transposable Element analysis

Transposable elements were putatively identified by searching the TAIR10 TE annotation against each genome separately using BLAST. Matches with an e-value < 1×10^−10^ were filtered out.

